# Prediction of RNA secondary structures in SARS-CoV-2 and comparison with contemporary predictions

**DOI:** 10.1101/2022.09.20.508790

**Authors:** Alison Ziesel, Hosna Jabbari

## Abstract

SARS-CoV-2, the causative agent of covid-19, is known to exhibit secondary structure in its 5’ and 3’ untranslated regions, along with the frameshifting stimulatory element situated between ORF1a and 1b. To identify further regions containing conserved structure, multiple sequence alignment with related coronaviruses was used as a starting point from which to apply a modified computational pipeline developed to identify non-coding RNA elements in vertebrate eukaryotes. Three different RNA structural prediction approaches were employed in this modified pipeline. Forty genomic regions deemed likely to harbour structure were identified, ten of which exhibited three-way consensus substructure predictions amongst our predictive utilities. Intracomparison of the pipeline’s predictive utilities, along with intercomparison with three previously published SARS-CoV-2 structural datasets, were performed. Limited agreement as to precise structure was observed, although different approaches appear to agree upon regions likely to contain structure in the viral genome.

## 1 Introduction

SARS-CoV-2 is the virus responsible for covid-19, the global pandemic that has infected more than 617 million people and killed more than 6.5 million people globally as of 18 September 2022 [1, 2]. It is a member of the clade *Betacoronavirus* and as such is a positive sense, single stranded RNA virus, with a genome size of 29,903 nucleotides. It has a likely zoonotic origin, with its most recent non-human host likely a bat species [3, 4]. Like other coronaviruses that afflict humans, including the previous SARS and MERS outbreaks, SARS-CoV-2 causes a primarily respiratory manifestation of symptoms, including dry cough, difficulty breathing and hypoxia, as well as non-respiratory symptoms including inflammation, thrombosis and myocarditis [5].

As an RNA virus, SARS-CoV-2 is capable of forming RNA secondary structure, which is the phenomenon by which an RNA molecule self-base pairs to form a non-linear structure. These structures then may carry out functions not specified directly by the genome sequence but rather by the topology of the formed structure. RNA secondary structure has been extensively characterized in viruses ranging from plant viruses to hepatitis C, HIV-1 and Zika virus [6, 7]. More specifically, coronaviruses are known to exhibit secondary structure; a great deal of work primarily in murine hepatic virus (MHV) and the alphacoronavirus 229E. In the case of SARS-CoV-2, there are three known domains capable of forming secondary structure: the 5’ untranslated region (UTR), the frameshift stimulatory element (FSE) and the 3’ UTR.

The 5’ UTR is predicted to contain at least five stem-loop structures, each of which are likely to play a role in the viral life cycle. Stem-loop 1 (SL1) is implicated in viral replication; experiments in the related virus MHV where this structure is disrupted lead to deficiencies in viral replication, which can be at least partially rescued by compensatory mutations [8]. SL1 is also involved in the mediation of long-range interactions with the 3’ UTR which permits replication of subgenomic RNAs (sgRNAs), an important feature of the coronaviral life cycle (reviewed in [9]). Stem-loop 2 (SL2) exhibits a highly conserved structure consisting of a five nucleotide pair stem and an exposed five nucleotide terminal loop containing a YUUGY sequence conserved across coronaviruses, hinting at a critical role [10]. SL2 is also implicated in the production of sgRNAs based on experimentation conducted in MHV and 229E [10, 11]. Stem-loop 3 (SL3) is observed in a subset of beta- and gammacoronaviruses, including SARS-CoV-2, SARS-Cov, and bovine coronavirus. In these viruses SL3 contains the transcription regulatory leader element (TRS-L) involved in the initiation of sgRNA synthesis; however in other betacoronaviruses including MHV this region is not folded. Selective 2’ hydroxyl acylation analyzed by primer extension (SHAPE) experimental data from MHV suggests that in those viruses where the TRS-L is contained within SL3, SL3 is a dynamic structure that may unfold to expose the TRS-L to its partner sequence TRS-B to facilitate sgRNA synthesis [12, 10]. Stem-loop 4 (SL4) consists of two stems separated by a small interior loop, referred to individually as SL4a and SL4b. SL4 is highly conserved across coronaviruses and while the complete deletion of SL4 in MHV is lethal, disruption of its structure is tolerated as is the deletion of either SL4a or SL4b [13]. This suggests SL4 may act as a functional spacer, meaning its physical presence is sufficient to enable 5’ UTR functionality. Stem-loop 5 (SL5) is a multibranch stem-loop structure with three terminal stem-loops, named SL5a through SL5c. While work in bovine coronavirus and MHV indicate that mutations of SL5a can impact viral replication, there is no consensus regarding the functions of SL5b or SL5c [14]. SL5 includes the start codon for ORF1a in its main stem, implying that this structure must be unfolded to initiate translation. In addition to these five SL structures known to exist in SARS-CoV-2, additional stem-loops are hypothesized based on computational analyses downstream of the start of ORF1a, although their biological significance and potential function remain uncharacterized [15].

The FSE occurs at SARS-CoV-2 genomic nucleotides 13462-13542 and is located in nsp12 at the junction between ORF1a and ORF1b. Structurally it consists of a seven nucleotide slippery site, a five nucleotide spacer, and a pseudoknot; when the processive ribosome encounters the pseudoknot, it retreats by one nucleotide within the slippery site, re-establishing the reading frame for translation and ensuring that ORF1b and downstream genes are read in the correct frame [16]. In SARS-CoV-2 there is some uncertainty as to the exact form the FSE pseudoknot takes as it appears to be dynamic (see [17] for an excellent summary) [18, 19]. The pseudoknot in conjunction with the slippery site stalls ribosome processivity, allowing slippage to occur, but only 15-30% of the time, thus maintaining a ratio of ORF1a:ORF1b where genes encoded by the polyprotein ORF1a are present in excess relative to those encoded by the polyprotein ORF1b [20, 21].

Lastly the 3’ UTR possesses a series of structures conserved in betacoronaviruses. Shortly after the stop codon of the N gene and overlapping with ORF10 is the structurally conserved bulged stem-loop structure (BSL) [22, 23]. Structurally the BSL consists of four stems interspersed with three unpaired bulge subsequences and terminated in a loop. Interestingly the base of the BSL is capable of interacting with an adjacent hairpin structure, forming a pseudoknot, where either the BSL first stem or the pseudoknot may be folded but not both. This arrangement is thought to act as a molecular toggle switch, oscillating between states to regulate functions pertaining to viral replication [24, 25]. The hairpin, known as P2, is part of a larger structural arrangement called the hypervariable region (HVR). The HVR exhibits very poor sequence conservation with the exception of an eight nucleotide sequence, and viruses with a complete deletion of the HVR are still replication competent *in vitro* [26]. Lastly, the 3’-most structure of the SARS-CoV-2 genome is the stem-loop II-like motif (s2m) element. The s2m exhibits an unusual tertiary conformation, resembling a portion of the *Escherichia coli* small ribosomal subunit RNA, leading to speculation that it may play a role in host translation interference [27, 28].

Since the onset of the current pandemic, a number of groups have leapt to the challenge of characterizing the secondary structure of the SARS-CoV-2 genome. Both computational predictive and molecular probing approaches have been pursued. In terms of biochemical probing, three key publications dominate: firstly, the Incarnato laboratory published their threefold SHAPE study, including both *in vivo* SHAPE and *in vitro* SHAPE and DMS probing describing the reactivities of nucleotides within the genome. They identified 87 regions of low reactivity, indicating strong likelihood of secondary structure formation [29]. However, they found that the known 3’ UTR structure did not meet their filtering criteria and the FSE structure was not correctly predicted. The second group to publish molecular probing data, the Pyle laboratory, also conducted SHAPE analysis but only *in vivo* [30]. They found stronger results for the 3’ UTR and FSE, and observed structure dispersed throughout ORF1ab and often abutting the junctions between coding elements. The third laboratory, Cliff Zhang’s, performed icSHAPE (in *vivo* click SHAPE) on infected liver cells and identified 37 structural elements; they also pursued analysis of other coronaviruses that infect humans and found similar structures in the 5’ UTR according to their protocol [31]. However, while they find excellent agreement with predicted 5’ UTR structures they do not identify a pseudoknot at the 3’ UTR, and make no mention of structure identified at the FSE.

Similarly, a number of computational biology studies have been undertaken for this problem. The Moss laboratory published their findings in early 2021, describing eight highly likely structures predicted with their ScanFold-based computational pipeline [32]. The Pyle laboratory, at nearly the same time, described their findings using the SuperFold RNA secondary structure prediction utility, identifying 61% of the genome as being base paired [33]. Additionally, the Das laboratory published their research into conserved sequences between SARS-CoV-2 and related coronaviruses using a combination of CONTRAfold and RNAz; they found 44 loci of predicted structure and identified components of the 5’ UTR, the FSE, and the 3’ UTR [34, 15]. Finally the Mathews research group applied a new iteration of their TurboFold folding algorithm, LinearTurboFold, to the complete SARS-CoV-2 genome and multiple sequence alignments, identifying 50 elements, 26 of which were not found in previous studies [35].

In contrast to what is already known regarding the UTRs and FSE, relatively little is definitively known regarding the capacity of structures to form in the central coding region of the SARS-CoV-2 genome. The possibility that SARS-CoV-2’s genome employs additional RNA structures to modulate its replicative cycle is appealing, as viruses must maximize their information content while maintaining a relatively compact genome. Use of unencoded structures overlaid on coding sequence, acting as regulatory elements, would be an efficient use of limited space, and is a tactic viruses already employ, the aforementioned FSE being a prime example. In a recent paper, Thiel and colleagues published details on a pipeline for identifying conserved ncRNA structures in vertebrate animals; similar work was also undertaken by Will et al [36, 37]. They make use of a number of existing computational utilities to find blocks of conserved sequence among a collection of vertebrate genomes, make an initial prediction regarding possible RNA structure, expand the subsequence containing the structure to reassess the propensity for structure formation, and then make a secondary prediction with the additional sequence information. This approach is very amenable to work with viral genomes as they are comparatively minute and densely informative, with very little genome space not devoted to protein encoding or regulatory function. The original Thiel pipeline as published could be employed with modest modifications to adjust for smaller, RNA-based genomes and to include examination for the possibility of covariation.

## 2 Materials and Methods

All informational resources used for this research are publicly available and free of charge. The genomes of various viruses, including the reference genome for SARS-CoV-2, were obtained from NCBI’s Nucleotide database [38]. All computational steps were performed using a Microsoft Azure Standard D16ds_v4 instance (16 vCPUs and 64 Gb of RAM), or a M1 MacBook Pro (16 Gb of RAM).

### 2.1 Modifications to original pipeline

As published, the Thiel et al. pipeline employed two utilities, RNAz and LocARNA [39, 40, 36]. In this work we have expanded upon those two utilities by the inclusion of the CaCoFold grammar-based structure prediction tool which enables the detection of both pseudoknotted base pairs and covariant nucleotide positions [41]. Further, the stringency of prediction performed by RNAz was increased from a threshold of 0.5 to 0.9. This equates to a change in specificity from 96% to 99%.

### 2.2 Virus selection

These analyses included viruses belonging to the genus *Betacoronoviridae,* coronaviruses known to infect humans, and closely related non-human host coronaviruses. Table 1 describes the host cell receptor protein, host organism, genus and sequence accession number of each virus studied.

**Table 1:** Viruses considered in this project. Receptors are angiotensin converting enzyme II (ACE2), dipeptidyl-peptidase 4 (DPP4), N-acetylneuraminic acid (Neu5Ac), alanine aminopeptidase (APN) and CEA cell adhesion molecule 1 (CEACAM1).

| Virus | Accession | Genus | Host | Receptor | Reference |
| --- | --- | --- | --- | --- | --- |
| SARS-CoV-2 | NC_045512 | betacoronavirus | human | ACE2 | [3] |
| SARS-CoV | NC_004718 | betacoronavirus | human | ACE2 | [42] |
| MERS-CoV | NC_019843 | betacoronavirus | human | DPP4 | [43] |
| HKU1 | NC_006577 | betacoronavirus | human | Neu5Ac | [44] |
| 229E | NC_002645 | alphacoronavirus | human | APN | [45] |
| NL63 | NC_005831 | alphacoronavirus | human | ACE2 | [46] |
| OC43 | NC_006213 | betacoronavirus | human | Neu5Ac | [47] |
| BCov | NC_003945 | betacoronavirus | cattle | Neu5Ac | [48] |
| HKU4 | NC_009019 | betacoronavirus | bats | DPP4 | [49] |
| HKU5 | NC_009020 | betacoronavirus | bats | unknown | [50] |
| HKU9 | NC_009021 | betacoronavirus | bats | unknown | [51] |
| MHV | NC_048217 | betacoronavirus | mice | CEACAM1 | [52] |
| RaTG13 | MN996532 | betacoronavirus | bats | ACE2 | [53] |

SARS-CoV-2, SARS-CoV1, MERS-CoV, HKU1, 229E, NL63 and OC43 were selected because they are all coronaviruses (albeit alphacoronaviruses in the cases of 229E and NL63) that infect humans (reviewed in [54]). BCov is evolutionarily related to OC43, with a fairly recent divergence between the two viruses. HKU4, HKU5, HKU9 and RaTG13 all infect bats, which may also be the origin of SARS-CoV-2; RaTG13 in particular is highly similar to SARS-CoV-2. Lastly the murine hepatitis virus MHV is similar to HKU1, as well as being a comparatively well-studied model virus.

### 2.3 Genome alignment

Whole genome alignment of all 13 viral genomes was performed using MAFFT and subsequently this preliminary alignment was used to construct a phylogenetic guide tree using unweighted pair group with arithmetic mean (UPGMA) hierarchical clustering implemented by MUSCLE [55, 56]. Construction of this phylogenetic tree is required to inform subsequent alignment by MULTIZ-TBA. MAFFT is a fast and accurate progressive alignment algorithm, making it well-suited to this purpose; MUSCLE is similarly fast and accurate, and is able to produce a phylogenetic tree from MAFFT output. Figure 1 shows the tree as calculated by MUSCLE. MULTIZ-TBA then produces aligned blocksets of sequence projected against a genome of choice, in this case SARS-CoV-2, using the previously generated guide tree [57]. MULTIZ-TBA performs all possible pairwise LASTZ local alignments between input sequences and then structures alignments into blocks representing conserved sequence between inputs. These conserved sequence blocks are then contextualized to emphasize conservation with the SARS-CoV-2 genome. Thus all aligned blocks of sequence obtained represent conserved regions of sequence in SARS-CoV-2 and their related sequences in one or more of the other viral genomes analyzed.

**Figure 1:**
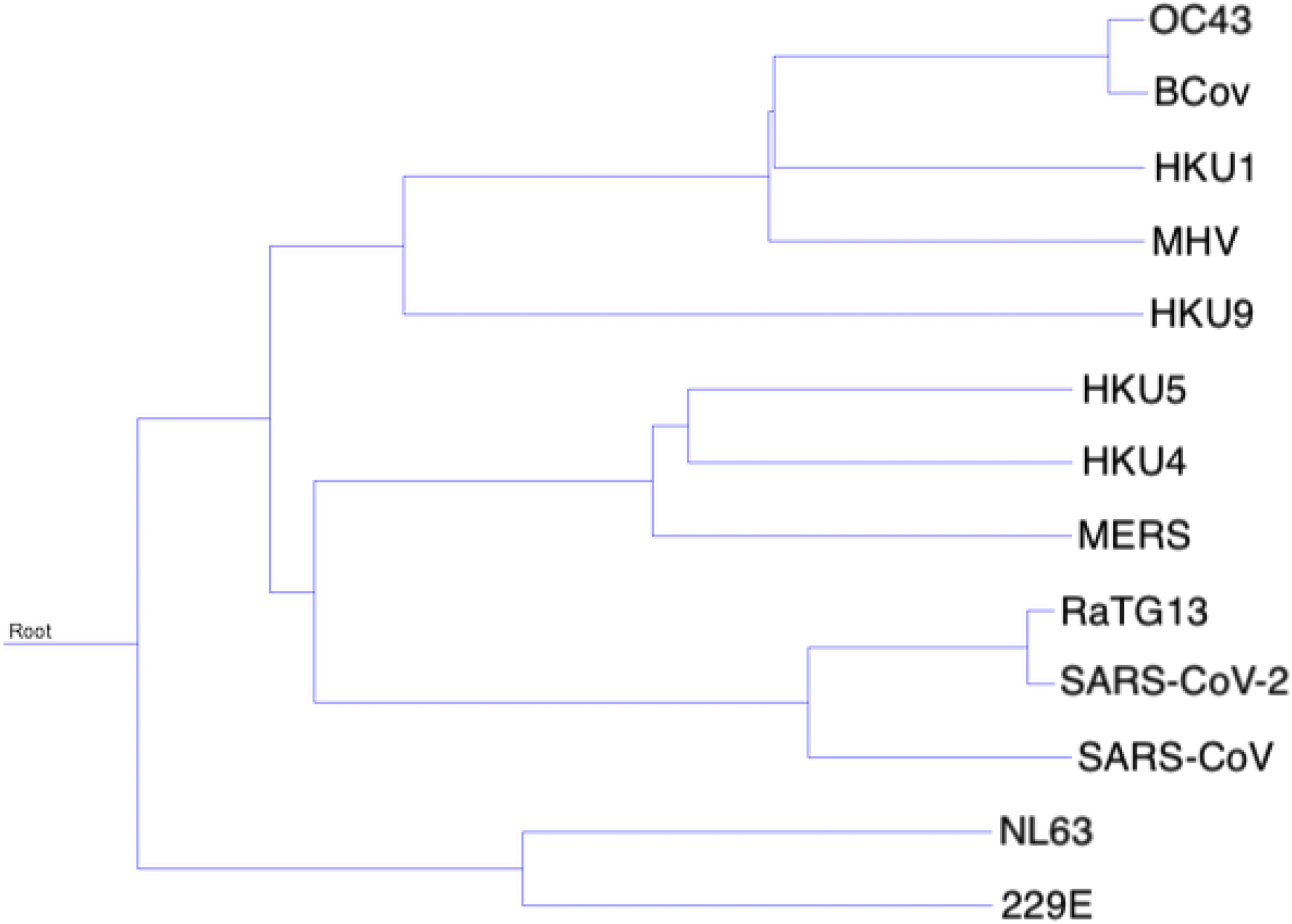
Phylogenetic tree describing the relationship between viruses as developed by MUSCLE from a MAFFT multiple sequence alignment.

### 2.4 Structure prediction

Following the pipeline outlined in [36], RNAz analysis was performed on all blocksets obtained from our earlier MULTIZ-TBA alignment. RNAz is a support vector machine (SVM) based predictor of RNA secondary structure and minimum free energy. Sequences that score very high for structural conservation and thermodynamic stability are more likely to form true RNA structures. Aligned blocks were prepared for RNAz analysis first by using the included rnazWindow.pl script, breaking longer blocks into 120 nucleotide windows with an overlap of 20 nucleotides. RNAz with default parameters was run on these resultant subblocks of sequence [58, 39].

A custom Perl script was developed to add 20 nucleotides of flanking genomic sequence to each sequence of each block of aligned sequence considered by RNAz, and these flanked sequences were then submitted to LocARNA for realignment using the arguments -probabilistic and -consistency-transformation; these arguments specify computation of reliability scores for predicted structures [40]. Flanked sequences were also resubmitted to RNAz for a second round of alignment again using default parameters.

To identify any covariant nucleotide positions the CaCoFold utility was also used on the flanked sequences [41]. CaCoFold uses positive and negative covariation data to identify sequence positions to be included and excluded from a folded structure respectively. CaCoFold is capable of predicting pseudoknotted structures (structures containing two substructures which intercalate), which RNAz is not intended for; however CaCoFold is sensitive to input sequence alignment quality, moreso than is RNAz. CaCoFold was run with default parameters. A summary of our modified pipeline can be seen in Figure 2. Note that for all three utilities, local structural predictions are made based on consensus sequence obtained from input multiple sequence alignment.

**Figure 2:**
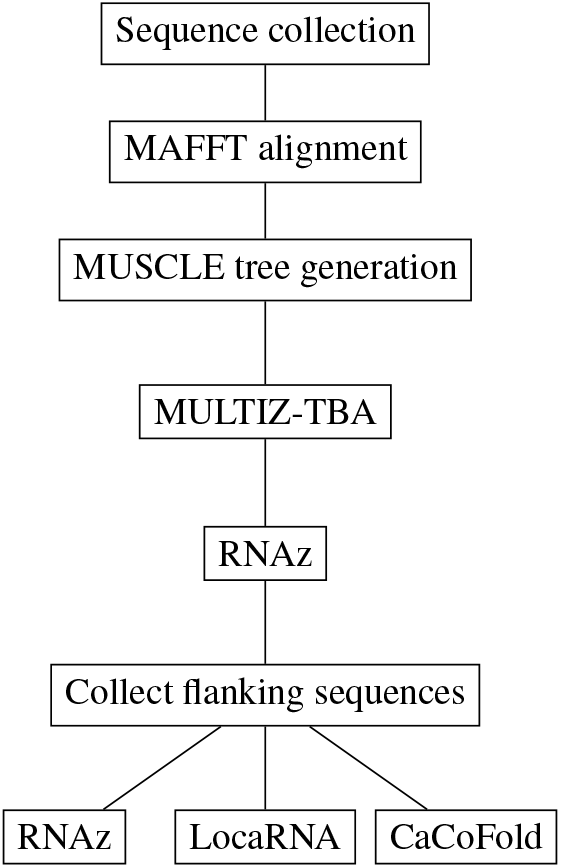
Modified pipeline employed in this study.

### 2.5 Comparison of predicted structures

Having generated predicted secondary structures using the RNAz, LocARNA and CaCoFold utilities, we next sought to identify commonly predicted elements among those three predictions. A Python utility was developed to compare dot bracket structures generated by each of the three predictions and identify common stems (regions of base pairing). Substrings within those structures that were agreed upon by all three predictive utilities that were capable of forming in the SARS-CoV-2 genome were identified for further analysis.

Additionally, distances between predicted structures were measured to quantify differences in predictions. Three distance metrics were chosen because they measure structural distances in slightly different ways. The base pair distance as measured by the ViennaRNA package is the number of pairs present in only one of two equal length dot-bracket structures. According to the documentation, this is “the number of base pairs that would have to be opened or closed to transform one structure into the other” ^1^. Hamming distance is the simple measure of how many characters in a string are non-identical at each position. In order to make this comparison, the unpaired bases (dots in a dot-bracket structure) were first converted to zeroes, while the paired bases (either left or right parentheses in a dot-bracket structure) were converted to ones. This decision was made on the basis that a nucleotide’s involvement in a base pair was more important than its identity as the 5’-most or 3’-most member of the pair. Lastly, Levenshtein edit distance was measured: it measures the number of substitutions, insertions, or deletions required to convert one sequence to another. As both DNA and RNA genomes may be changed by insertions, deletions, or point mutations, it was believed that this may be an appropriate representation of structural distances as well. Python tools to calculate the Hamming distance and Levenshtein string edit distance were developed and these distance metrics, along with the ViennaRNA bp_distance metric were collected [59].

### 2.6 Comparison with published findings

Three published reports describing potential genomic RNA secondary structures in SARS-CoV-2 were considered. Firstly the paper published by Andrews et al., henceforth referred to as the Andrews data set, describes an undertaking to predict RNA secondary structures using a computational biology pipeline [32]. Secondly works by Huston et al. and Manfredonia et al. detail biochemical analyses of the secondary structures observable in the SARS-CoV-2 genome, and are referred to by their first authors throughout this work [29, 30]. The Huston data set is derived from *in vivo* observations using SHAPE with 200mM 2-methylnicotinic acid imidazolide (NAI) as the SHAPE reagent, and were analyzed using ShapeMapper2 [60]. The Manfredonia data set includes both *in vivo* and *in vitro* SHAPE data using 100mM NAI, as well as using 150mM dimethyl sulfate (DMS) *in vitro.* The Manfredonia data set was analyzed using RNA Framework v2.6.9 [61].

For each of the three data sets, genomic regions corresponding to those predicted as significantly likely to produce RNA secondary structure in this work were identified and the dot bracket RNA secondary structure predictions from each publication were collected. In the case of the Andrews data set a Python utility was written to convert their .bp formatted data to dot bracket results. Note that in the Andrews, Huston and Manfredonia data these structures represent globally generated structures and as such, the genomic subregions corresponding to local predictions produced by this pipeline include ‘broken’ structures with incomplete base pairs cut off by the window size specified. Levenshtein distances were measured between the structures predicted in this work and between each pair of published results, generating a four way pairwise distance matrix.

## 3 Results

### 3.1 Prediction of secondary structures

Following our pipeline, 40 blocks of aligned sequence were found to have high confidence predictions of conserved RNA secondary structure, summarized in Figure 3. High confidence predictions required at least two species with an RNA class probability score greater than 0.9; this was a modification from the original pipeline, increasing specificity. Interestingly a number of these significant structures appear near the junction between protein encoding regions in the polyprotein transcript ORF1ab especially between nsp1/2 and nsp2/3 with conservation among SARS-CoV-2, SARS-CoV and RaTG13, and between other genes, notably ORF6/ORF7a, ORF7b/ORF8, and between the envelope and M genes. Much of ORF3a, which encodes a transmembrane protein involved in replication and viral release from the host cell, is predicted to have high confidence RNA secondary structure, again within the trio of viruses SARS-CoV-2, SARS-CoV and RaTG13.

**Figure 3:**
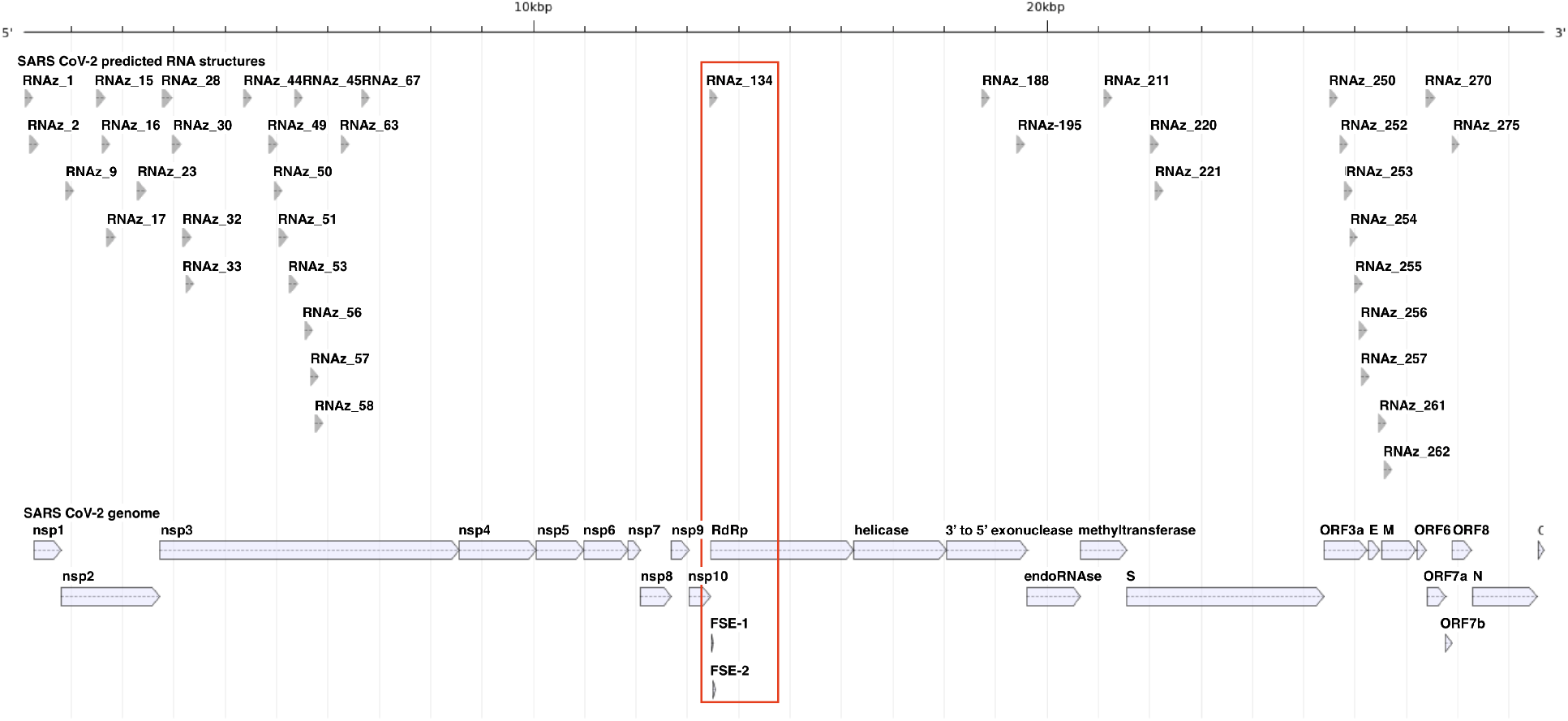
Positions of the 40 likely RNA structures (top) aligned against the SARS-CoV-2 genome (bottom). The location of the sequence block with both high confidence RNA structure and covariant bases, which overlaps with the FSE, is indicated with a red box.

Following the modified pipeline, we undertook to predict possible pseudoknotted structures as well as identify covariant nucleotides, which are indicative of conserved structure despite apparent sequence divergence. Pseudoknots are known to be functionally relevant to virus biology, as in the case of the FSE. However, no pseudoknots were predicted by CaCoFold in any of the 40 structures indicated to be valid by RNAz. 40 blocks exhibited covariant nucleotide pairs, with limited overlap with those high confidence RNA structure predictions as detected by RNAz: only one block, that spanning the FSE, was both high confidence and exhibited covariant bases (Figure 3). Covariant bases external to likely RNA structures within the coding region of SARS-CoV-2 may still represent points of interest for viral function albeit not in RNA structures. Structures RNAz_0 and RNAz_1 localize entirely within the 5’ UTR while RNAz_2 corresponds to the 3’ terminal of the 5’ UTR and the beginning of the coding region of nsp1, spanning genomic nucleotides 179-338. Rfam, a publicly available repository of RNA secondary structures, contains entry RF03120 ^2^, which includes a predicted structure of the 5’ UTR of sarbecoviruses, a subgenus of the *Betacoronovirus* to which SARS-CoV-2 and other SARS-like viruses belong. While RF03120 is based on comparative sequence analyses, it is generally accepted that coronavirus 5’ UTRs contain a series of stem-loop structures, with a number of studies thoroughly reviewed by [62]. RNAz_0, starting at the beginning of the genome, overlaps with stem-loops 1 through 4. RNAz_1 overlaps with SL4 and a portion of 5, while RNAz_2 overlaps with stem-loop 5 (SL5) of the sarbecovirus predicted 5’ UTR. As each of the blocks of sequence cover a subgenomic window rather than encompassing the entire 5’ UTR sequence, RNAz_1 and RNAz_2 do not fully recapitulate the structure of SL5 but it does correctly identify that genomic region as capable of forming RNA secondary structure. As it begins from the beginning of the genome RNAz_0 performs quite admirably in predicting the consensus structure. When considering only whether a base is involved in a base pair, the calculated Levenshtein distance between the genomic window for sarbecoviruses and RNAz_0 is 17, RNAz_1 is 60, and RNAz_2 is 51. Structure RNAz_134, the only structural element predicted to have covariant nucleotides and high confidence structure, spans the SARS-CoV-2 frameshifting stimulatory element (FSE), a critical genomic element responsible for correct translation of downstream genes located nucleotides 13462-13542. RNAz_134 spans nucleotides 13314 through 13470, including upstream sequence and the first six nucleotides comprising the FSE. While Rfam entry RF03120 includes a predicted FSE consisting of a single stem and a pseudoknot, the work of Zhang and colleagues has demonstrated through cryo-electron microscopy that the SARS-CoV-2 FSE consists of a three stem pseudoknot [63]. Of the three predictive tools used in this study, only CaCoFold is capable of predicting pseudoknots, but in this case it did not predict a pseudoknot in the genomic window identified by RNAz as likely to harbour RNA secondary structure. CaCoFold did predict one pair of nucleotides in this genomic window as being covariant: window nucleotides 43 and 141, which correspond to genomic nucleotides 13357 and 13455, both of which are upstream of the identified FSE.

The remainder of the forty identified likely structures did not overlap with any further known RNA secondary structures in the SARS-CoV-2 genome and were instead within coding regions. Five structures are contained within nsp2, 15 within nsp3, 2 within nsp14, 3 within nsp16, and 7 within ORF3a. Four structures spanned the boundaries between coding units. RNAz_261 and RNAz_262 span the boundary between E and M, RNAz_270 between ORF6 and ORF7a, and RNAz_275 spans the ORF7b/ORF8 boundary. Supplementary table 2 details the structures predicted by each of the three structure prediction utilities.

### 3.2 Comparison of predicted structures

Of those 40 structures predicted as likely to represent true RNA secondary structure, ten of them exhibited substructures predicted in common by all three predictive utilities, RNAz, LocARNA and CaCoFold. In no cases was the complete local structure predicted by all three utilities, rather, portions of the structure, with internal stems and terminal stem loops were most likely to be agreed upon. Figure 4 shows those ten structures with their three-way mutual common elements indicated. Structure RNAz_2 spans the end of the 5’ UTR and into the 5’ end of ORF1a/nsp1. Structures RNAz_9, 15 and 17 are located in ORF1a/nsp2, while 28, 30, 32, 33 and 54 map to ORF1a/nsp3. Structure RNAz_254 is found in ORF3a.

**Figure 4:**
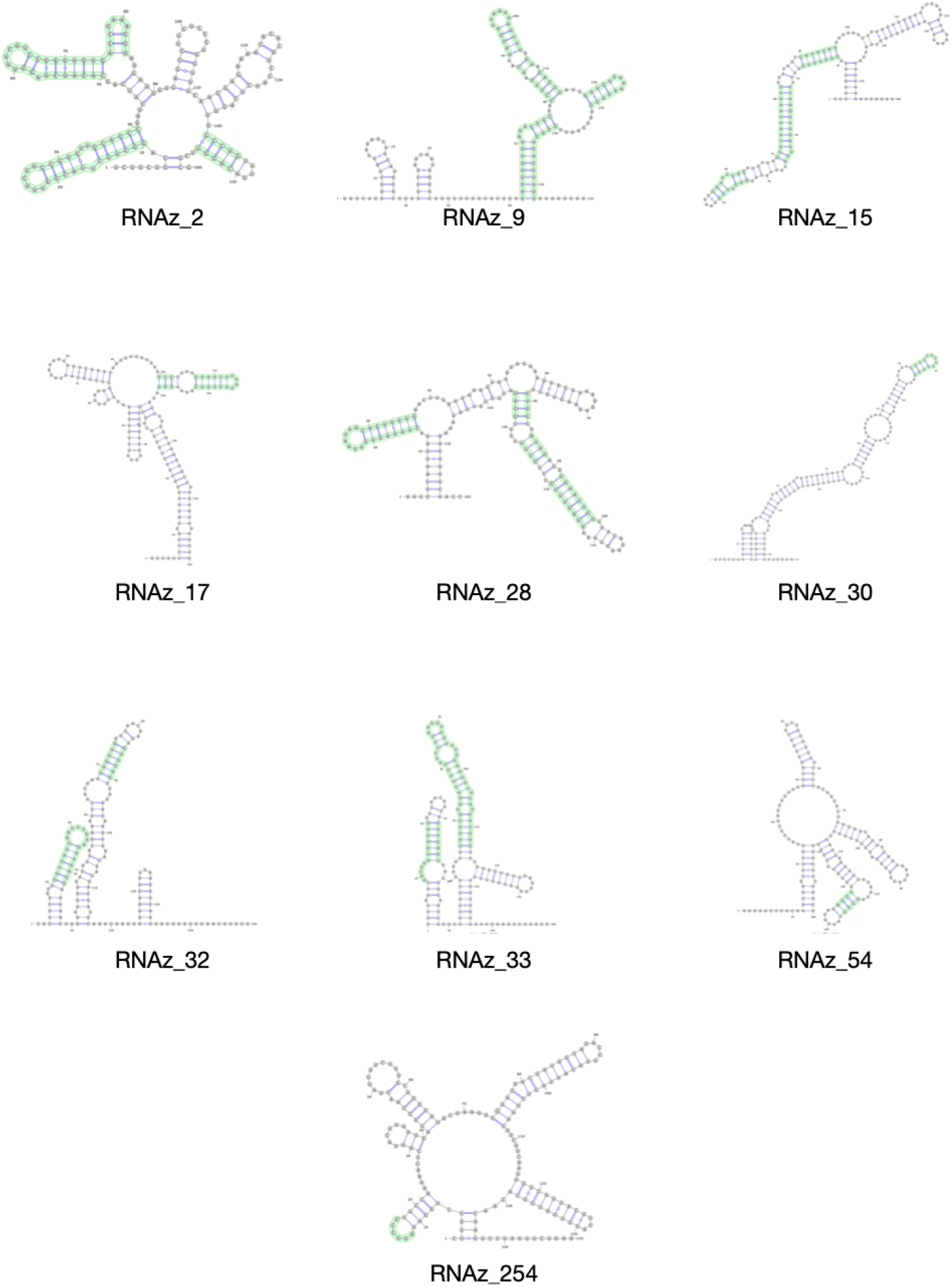
Structures of the 10 predicted RNA structures with agreed upon elements highlighted in green.

In addition to identifying commonly predicted subsequences, the differences between each prediction for each of the 40 structures were quantified by measuring the Hamming distance for bases involved in base pairing, the Levenshtein string edit distance, and the base pair distance as implemented by ViennaRNA [59]. Recall that for each of RNAz, LocARNA and CaCoFold the obtained structures are consensus structures developed from two or more of the viruses aligned for a given blockset amongst the total virus set used in this study. Supplementary table 3 describes the distances obtained from each trio of predicted structures for each blockset.

### 3.3 Comparison of results with published findings

In order to quantify the distances between structures predicted by the Andrews, Huston and Manfredonia papers and this work, the Levenshtein edit distance was calculated between each pair of structures and a pairwise distance matrix was developed for each of the 40 likely structures identified here; the distance matrices can be seen in Figures 5 and 6. The RNA secondary structures predicted by RNAz were used as the representative structures for this work. Four major patterns of distance emerged: (1) all four structures were relatively close to one another, indicating similarity of prediction; (2) all four structures were relatively distant, indicating general disagreement as to bases involved in stem formation; (3) the SHAPE data sets (Huston and Manfredonia) were relatively close to one another and relatively distant from the Andrews data and this work, and (4) the Andrews data set was relatively distant from the other three data sets. Examples of these patterns include RNAz_9, 16 and 23 for pattern (1), pattern (2) is exhibited in RNAz_44, 53 and 220, pattern (3) can be seen in RNAz_30, 250 and 252, and RNAz_1, 15 and 221 are examples of pattern (4). In addition to the four observed patterns of distance, some exceptional distance matrices emerge. RNAz_0 shows remarkably short distances between all four structures, but as it is the first possible structure that might be formed, this may be due to the fact that there is no upstream sequence that could form larger structures in the SHAPE data sets. RNAz_2 shows modest agreement between my work and the SHAPE data of Huston and Manfredonia, but the predicted Andrews data is extremely different. RNAz_250 is interesting in that the two predicted structures, from this work and the Andrews paper, are quite far apart while being moderately distant from the two SHAPE structures, implying a divergence in how the two predictive pipelines examined this region. The overlapping structures RNAz_254, 255, 256 and 257 all have quite short distances between structures, indicating a consensus regarding the state of this portion of the SARS-CoV-2 genome, which is within the ORF3a gene. In terms of overall patterns across all forty structures, the SHAPE data sets of Huston and Manfredonia had the shortest overall distance with a mean distance of 33.225, followed by the distance between this work and the Manfredonia data set with a mean distance of 44.0. The greatest observed distance was between this work and the predictive data set of Andrews, measuring a mean distance of 53.35. Overall the SHAPE data sets most resembled one another, then the predictive and SHAPE data sets resembled one another intermediately, with the greatest distance between the two predictive approaches.

**Figure 5:**
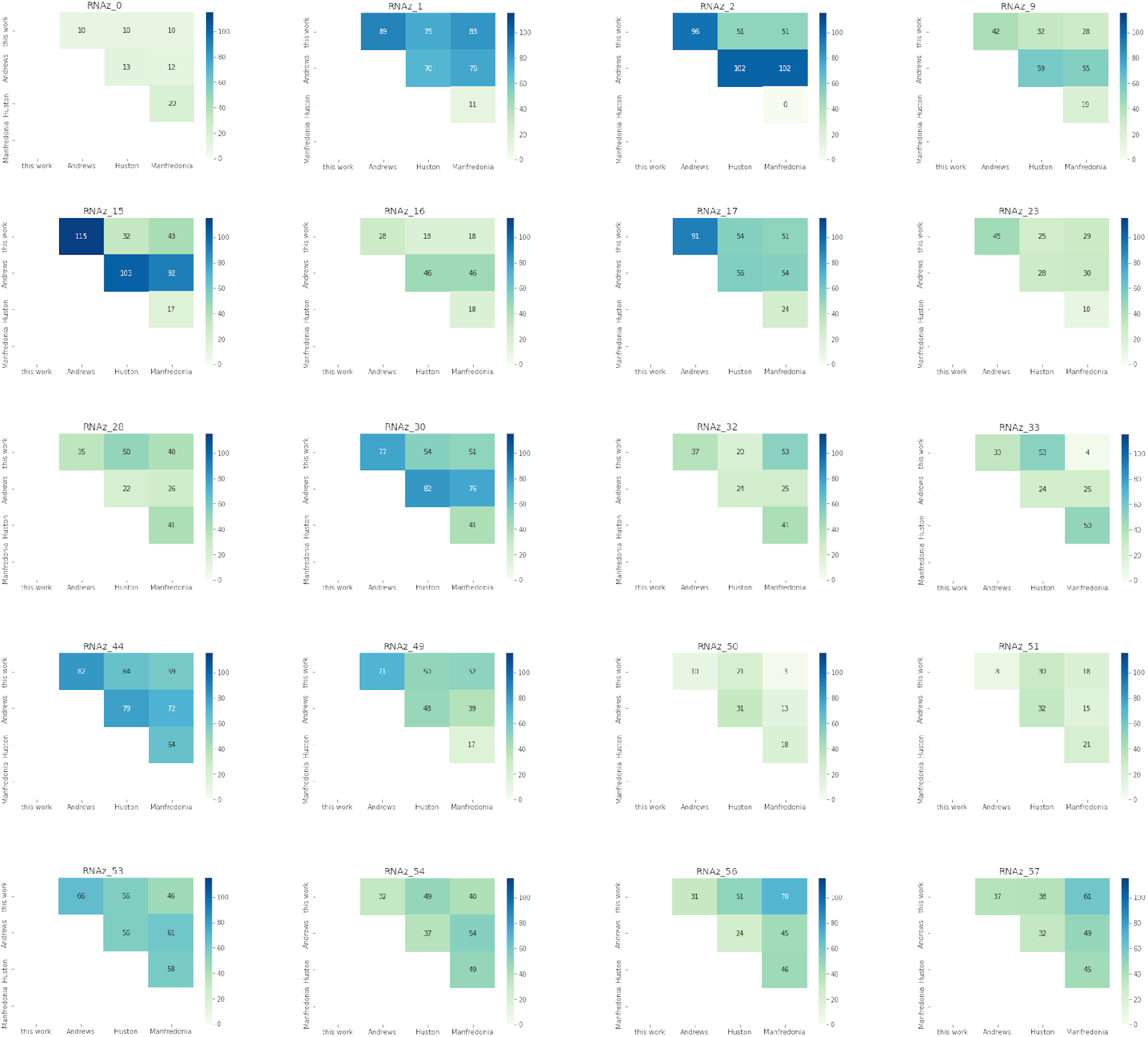
Part one of the distance matrices describing edit distance between this work, the predictive work of Andrews, and the SHAPE data of the Huston and Manfredonia papers.

**Figure 6:**
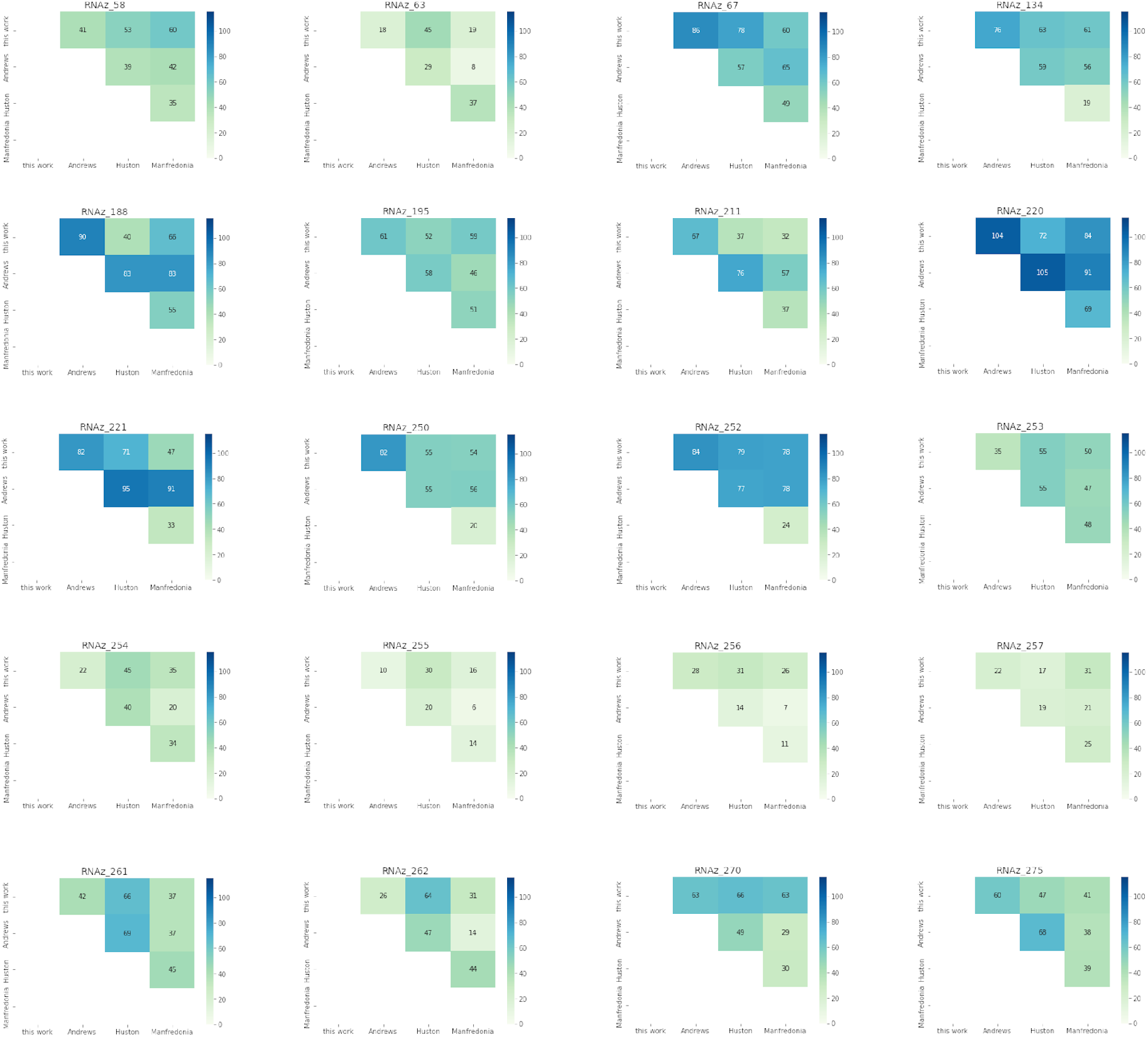
Part two of the distance matrices describing edit distance between this work, the predictive work of Andrews, and the SHAPE data of the Huston and Manfredonia papers.

In order to identify statistically significant differences in distances between pairs of predictions, one-way analysis of variance was performed using the Kruskal-Wallis test; this test was chosen as the distribution of differences in distances was not normally distributed in all cases. The result of the Kruskal-Wallis test was an H-statistic of 16.76687, with a p-value of 0.00496, rejecting the null hypothesis at alpha equal to 0.05. This was followed up with Dunn’s post hoc tests to identify which set of distances were statistically significant. The difference in distances between Huston/Manfredonia and Ziesel/Andrews, Ziesel/Huston and Andrews/Huston were statistically significant with p-values of 0.00996, 0.04538 and 0.00617 respectively.

## 4 Discussion

Forty subregions of the SARS-CoV-2 genome were predicted by our pipeline as very likely to contain RNA secondary structure. These structures tend towards the 5’ end and 3’ third of the genome and cover structure already known to exist in the SARS-CoV-2 genome, including portions of the 5’ UTR and FSE, although the structures predicted here do not perfectly recapitulate those putative structures. Following viral selection, multiple sequence alignment, and identification of conserved contiguous sequence, these 40 regions were selected first based on their high levels of conservation with related coronaviruses, and secondly due to their relatively high levels of structural conservation and thermodynamic stability as calculated by RNAz. In this work we employed an increased RNA class probability score to maximize the specificity of prediction of structure-containing sequences. This choice has surely resulted in a number of false negative predictions; additionally, the use of relatively small windows for analysis (120 nucleotides in the first round, and 160 nucleotides for the flanked sequences in the second round of analysis) may artificially obscure larger structure by focusing on too small subregions. Further analysis of these regions was performed following a modest expansion of the regions of interest and then reanalysis using RNAz and the additional RNA structural prediction utilities LocARNA and CaCoFold. The expansion of regions of interest with an additional twenty nucleotides on either side of the identified conserved regions is done so as to better capture a potential structure by including its less well-conserved sequence that may nonetheless be involved in formation of a conserved structure. Reanalysis with RNAz in this larger subgenomic context, again considering the structural conservation of predicted structure, along with its thermodynamic properties captures any potential larger structure.

This is followed by the LocARNA structural prediction utility, which was originally developed as a companion to RNAz [40]. LocARNA considers the problem of likely RNA secondary structure with a different algorithmic approach: RNAz first performs alignment of the input multiple sequence data before predicting a structure, but LocARNA performs alignment and structural prediction simultaneously, using information from each sub-problem to inform the other. In particular, LocARNA claims structural boundary prediction as a strength, due to the calculation of sequence-structure-based alignment reliability (STAR) scores for each nucleotide in an alignment. The reliability score assesses the likelihood that each nucleotide in a potential structure contributes to the final structure and so can be used to limit false positive base pairs in a structural prediction. In this study we did not find that boundary definition as determined by LocARNA was especially informative; this may be due to the small window size previously mentioned, or it may be that the initial steps of multiple sequence alignment and identification of conserved sequence blocks generated natural boundaries of structure. Another possibility is that, unlike in vertebrate genomes and ncRNAs, nearly all of the SARS-CoV-2 genome has the additional constraint of encoding viral proteins as well as specification of any potential structure. While the addition of flanking, less-conserved sequence may have partially alleviated the artificiality imposed by analyzing conserved sequence blocks, it may not have been sufficient for LocARNA to perform as well as it did in the originally published Thiel pipeline.

Lastly we included the CaCoFold structural prediction utility, a novel addition to the original Thiel pipeline. Like RNAz and LocARNA, CaCoFold operates on a multiple sequence input, but it does not seek to minimize thermodynamic parameters; rather it applies stochastic context free grammar (SCFG) rules to generate a structure for a consensus alignment. CaCoFold also calculates covariation statistics to identify pairs of covariant nucleotides, which can provide powerful evidence for positions highly important to structure formation. Very few covariant nucleotide pairs were identified among the viral genomes used in this study, and only one pair was identified in a sequence that met our threshold for structure likelihood according to RNAz. As a result, further characterization of covariant nucleotide pairs was not pursued in this work. It is possible that insufficiently variant alignments may suppress the detection of covariant positions, and as a result of using conserved sequence blocks for analysis with CaCoFold, the covariation potential may have been minimized [41]. This may be ameliorated either by the inclusion of more and more diverse viral genomes in the initial stages of the pipeline, or the adjustment of the MULTIZ-TBA alignment parameters to be more tolerant of nucleotide-level differences in a larger block of conserved sequence.

A number of structures predicted are near to or span the junction between viral genes, inviting the speculation that they may represent flow control elements to regulate expression of downstream genes. Four structures are predicted to span the boundaries between coding regions: two overlapping regions of structure span the E/M gene boundary, one spans the ORF6/ORF7a boundary, and one spans the ORF7b/ORF8 boundary. Moreover, this observation is echoed in the work of Huston et al.: they also observe structure at or near the same locations [30]. Further investigation into the potential function of these regions, including possible RNA binding protein interacting partners, is warranted. Additionally, much of the coding sequence of ORF3a appears to be capable of forming RNA secondary structure. The significance of this is not yet clear and also deserves closer scrutiny.

Of the forty predicted regions likely to contain secondary structure, ten were identified as having substructures commonly predicted by RNAz, LocARNA, and CaCoFold. As described above, each of these three utilities takes a different algorithmic approach to the prediction of secondary structure, and the convergence of predictions may indicate increased reliability of the prediction for those substructures. Indeed, the approach taken in this work was deliberately developed from the previously described pipeline to incorporate additional algorithmic glimpses at possible genomic structure. A single methodology is an inherently limited view of the data and by employing a broader approach with three different algorithms better enables a more complete view. Of the ten commonly predicted substructures, one element is located in ORF3a, while the majority are located in ORF1a, specifically in nsp1, nsp2, and nsp3, including one substructure that spans the 5’ UTR/ORF1a boundary. This region has been predicted to contain additional stem loops that may be functionally associated with the 5’ UTR but are present in coding sequence [15].

Beyond the commonly predicted substructures, modest agreement between the three structural predictions for each structure was observed. The mean Levenshtein distance observed between RNAz and LocARNA predictions was 59.95, between RNAz and CaCoFold was 67.25, and between LocARNA and CaCoFold was 54.2. It was expected that the two thermodynamic-based utilities, RNAz and LocARNA, would have the most similar predictions but this was not the case: LocARNA and CaCoFold showed approximately 10% reduced distance relative to RNAz and LocARNA. It is reasonable to speculate that the grammar rules applied by CaCoFold prioritize similar features to LocARNA’s aim to identify strong structural element boundaries. Regardless of the nature of the differences and similarities between the three approaches, there is no clear consensus structure prediction for the majority of structures.

This lack of consensus extends to other data sets produced employing different approaches. The previously described Andrews data set uses only the reference sequence for SARS-CoV-2, eschewing multiple sequence input to focus strictly on the single genome. They conduct their analysis using the SuperFold pipeline. Both the Huston and Manfredonia SHAPE analyses employ NAI as their SHAPE reagent, but they use different computational approaches to analyze their results: the Huston data set uses ShapeMapper2, while the Manfredonia data was analyzed using RNA Framework. As with the Andrews data set, the SHAPE studies considered here focus specifically on the single SARS-CoV-2 reference genome. As described in the results, the SHAPE-predicted structures had the shortest distance between predictions with a mean Levenshtein distance of 33.225, with the greatest distances observed between this work and the Andrews computationally-predicted structures. A statistically significant difference was found in the distances between the smallest and greatest observed mean distances. While it is not surprising that the SHAPE-based approaches are most similar, they still differ appreciably, despite using the same SHAPE reagent (albeit at different concentrations). Whether their differences are due to laboratory handling or to the different analytical pipelines employed is unclear. In terms of the difference in distances between the computational Andrews data and the original computational predictions in this work, a great deal of difference is likely due to the fact that this analysis is rooted in a multiple sequence alignment of related coronaviruses, as opposed to the single SARS-CoV-2 reference genome sequence. In this work, we focus on structure predicted in sequence conserved amongst the thirteen chosen coronaviruses used in the multiple sequence alignment and so will not detect structure in sequence recently evolved in SARS-CoV-2. That said, a number of the structures predicted in this work were based on a sequence alignment between the triad of SARS-CoV-2, SARS-CoV and RaTG13, a bat-hosted betacoronavirus that shares 96.1% nucleotide sequence identity with SARS-CoV-2 [64]. The similarity between SARS-CoV and SARS-CoV-2 is 82.5% [65, 66]. The most recent common ancestor of RaTG13 and SARS-CoV-2 may have existed as little as fifty years ago [67]. Any SARS-CoV-2 specific sequence not also present in RaTG13 would represent a very recent acquisition indeed and may not yet have concretized into a functional structure.

This study makes use of the Levenshtein string edit distance to quantify the distance between RNA secondary structures are represented by dot bracket notation, both within the original work presented here and between this work and previously published data sets. Informally, Levenshtein distance is the number of single character changes required to convert one string into another, including insertions, deletions, and alterations of individual characters, which are analogous to those changes that might take place in a genomic context. We have chosen to focus on the dot bracket notation representation of secondary structures rather than primary genomic sequence as structure is not unique to sequence, and so divergent sequences may nonetheless specify the same RNA secondary structure. Treating the structure as a one dimensional string representation greatly facilitates quantifiable comparisons between candidate structures, and Levenshtein string edit distance is only one measure applicable: competing measurement systems include Hamming distance (the number of dissimilar positions in two strings of equal length) and the ViennaRNA implementation of base pair distance (the number of base pairs present in one structure but not the other). Hamming distance was deemed too simplistic for this study as a one base pair insertion or deletion would result in a very large Hamming distance despite possibly not interrupting the greater genomic structure. Base pair distance was not employed because it presupposes the two structures are of equal length, which was not always a valid precondition in the comparisons undertaken in this study. As any RNA secondary structure may be easily represented in dot bracket notation, the application of Levenshtein distance as a metric for structural dissimilarity is easily extensible to any number of comparisons with other SARS-CoV-2 or related viral structural predictions.

## 5 Conclusions

Modifying and repurposing an existing computational pipeline designed for the detection of ncRNAs in vertebrates has captured three different estimates of RNA secondary structure in regions of the SARS-CoV-2 genome that is conserved in other coronaviruses, including those that infect humans and bats. We have chosen different structural prediction algorithms to leverage their distinct strengths, and we have relied on a multiple sequence alignment input to capture those structure-harbouring regions most likely to be functionally relevant. We have compared our predictions with one another, characterizing them in terms of their commonly predicted substructures. Further, we have compared our findings with three previously published data sets, both computational and biochemical in nature. This demonstrates that there is no consensus structure among any of the predictions considered, and we believe this indicates that each approach is likely identifying regions that can generate structure but may not be adequately assessing that structure. It is possible that in some cases, we and the other data sets are capturing an average of an ensemble of multiple possible structures. Further work is required to elaborate upon this possibility. While there is no consensus prediction, these studies are important as they can inform downstream analyses, refining their focus to the most likely structural possibilities. Ongoing research into the secondary structure of the SARS-CoV-2 genome is still required.

---

**Supplementary Table 1:**
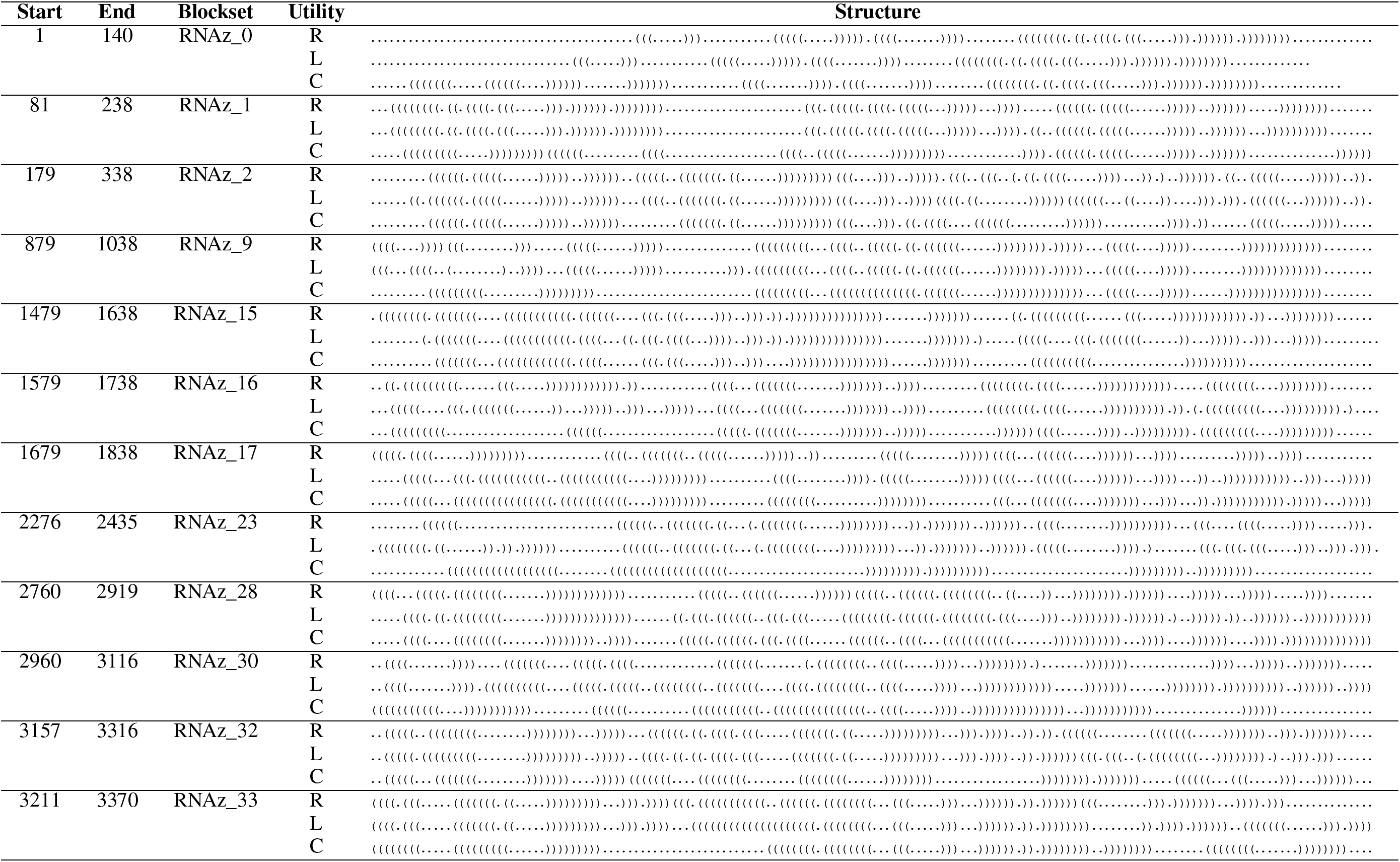
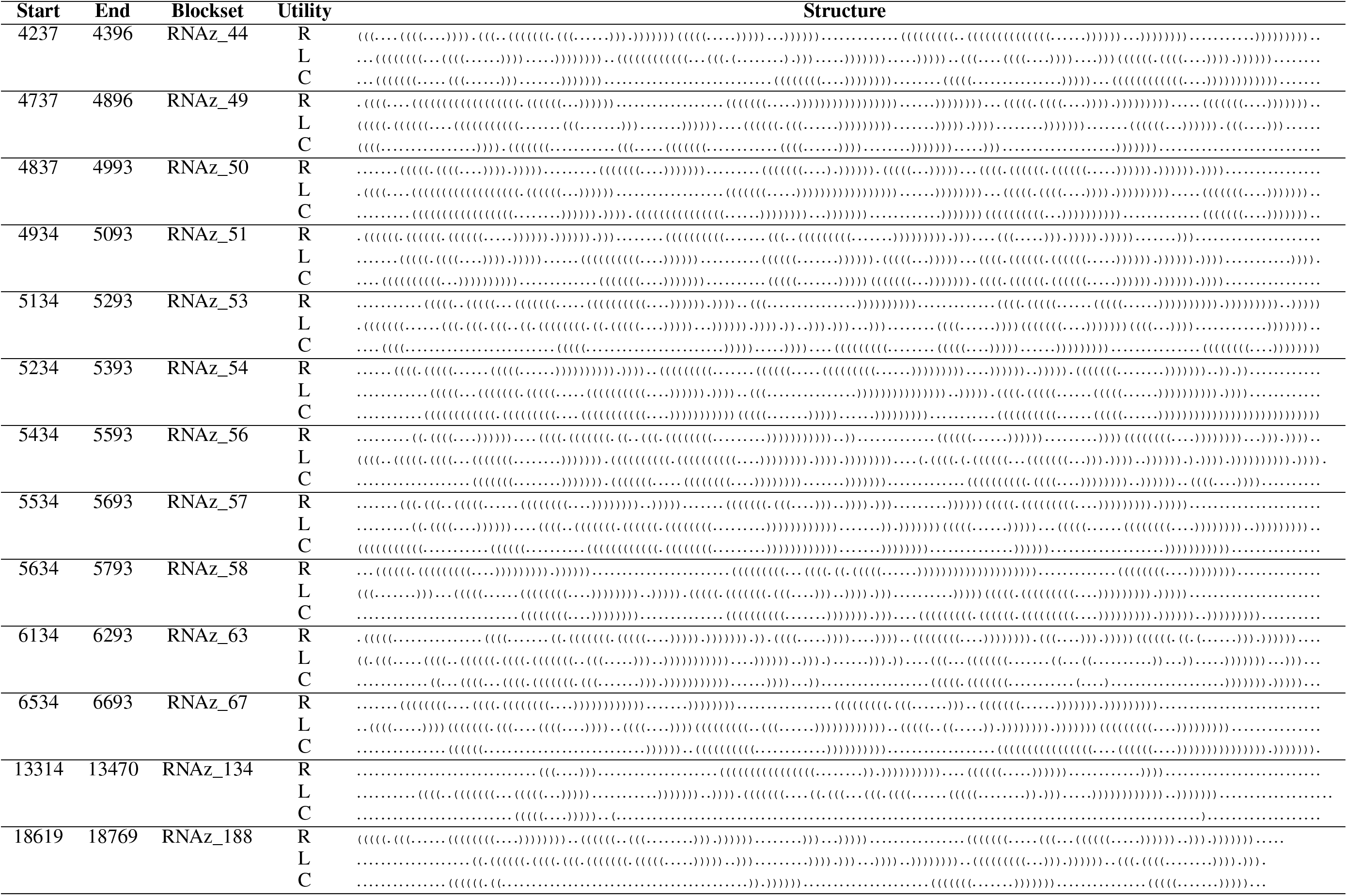
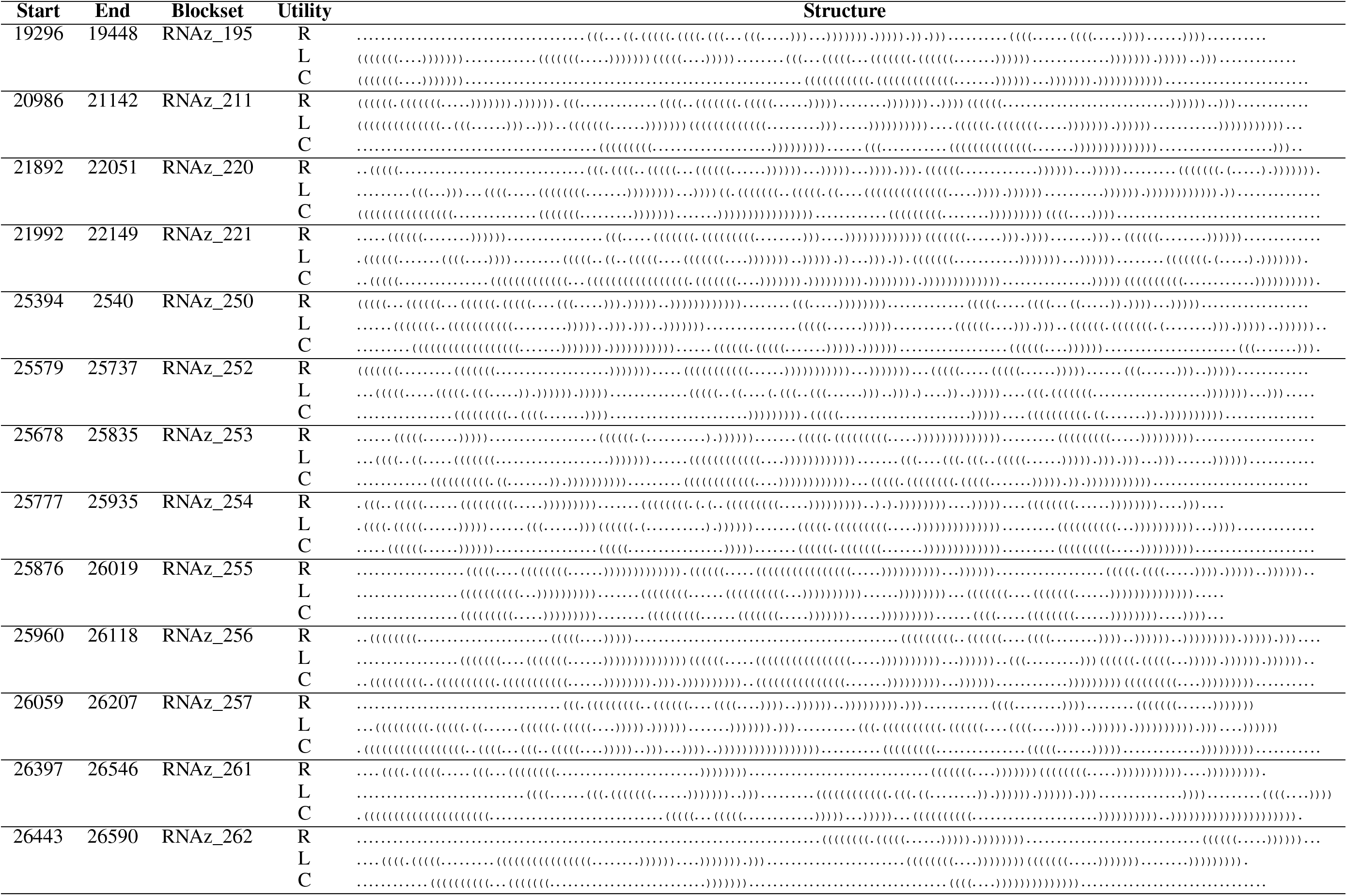
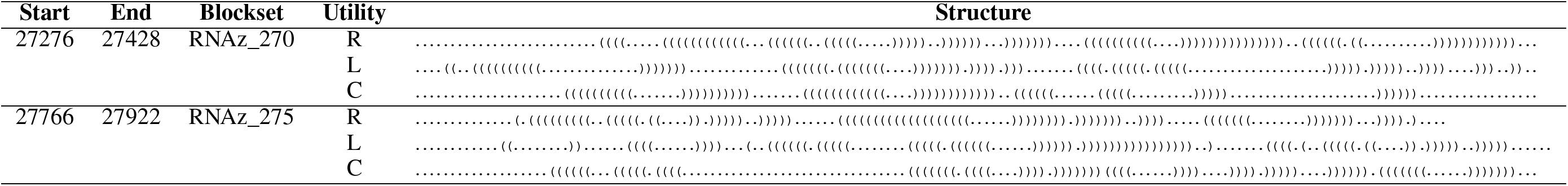
Predicted structures, and the SARS-CoV-2 location of each blockset. ‘R’ is RNAz, ‘L’ is LocARNA, ‘C’ is CaCoFold.

**Supplementary Table 2:**
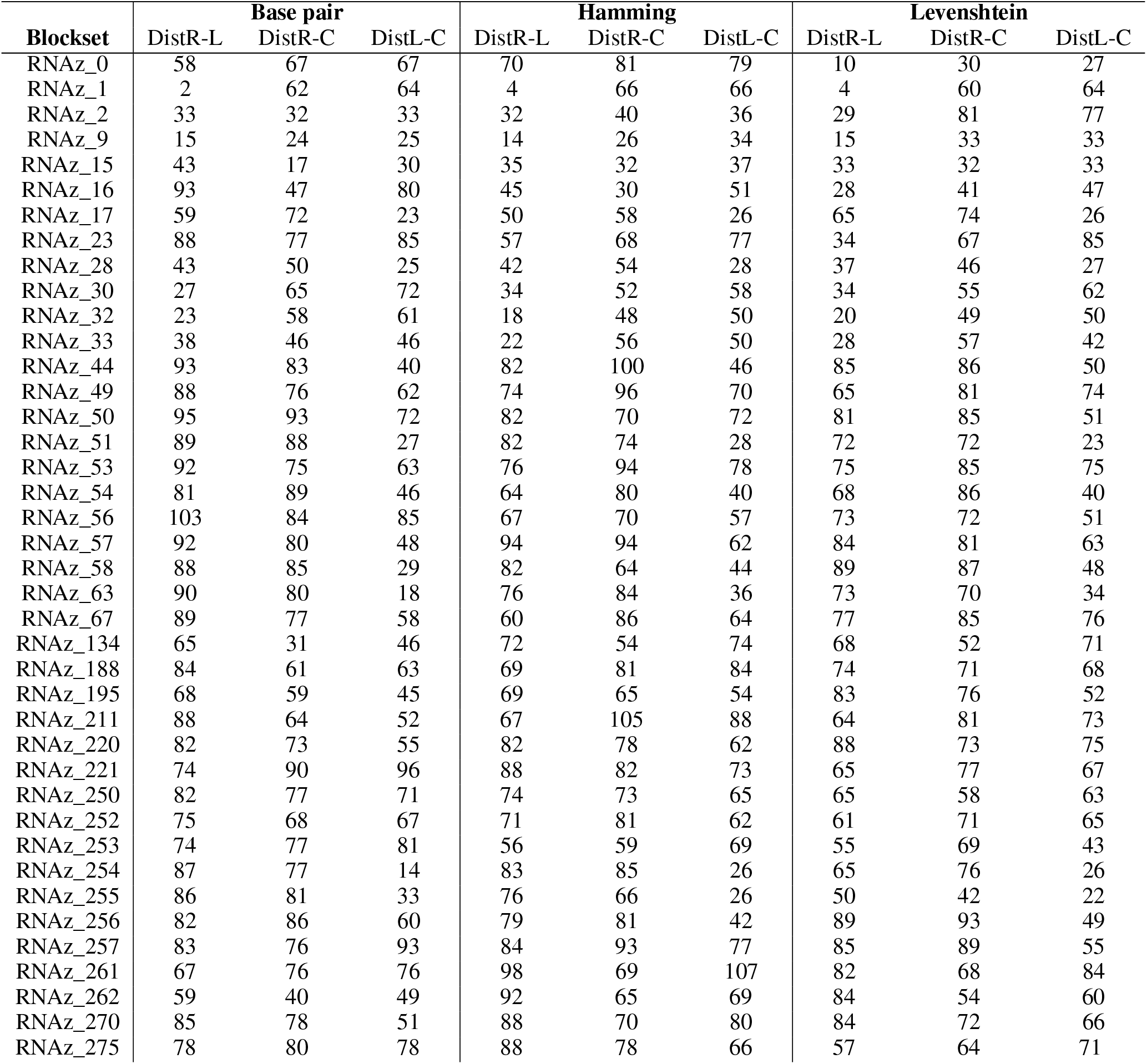
Base pair, Hamming, and Levenshtein distances between predicted structures.

## Footnotes

1 https://www.tbi.univie.ac.at/RNA/ViennaRNA/doc/html/distance_measures.html

2 http://rfam.xfam.org/family/RF03120

## Notes

### Competing Interest Statement

The authors have declared no competing interest.

